# Statistical learning in epilepsy: Behavioral, anatomical, and causal mechanisms in the human brain

**DOI:** 10.1101/2023.04.25.538321

**Authors:** Ayman Aljishi, Brynn E. Sherman, David M. Huberdeau, Sami Obaid, Adithya Sivaraju, Nicholas B. Turk-Browne, Eyiyemisi C. Damisah

## Abstract

Statistical learning, the fundamental cognitive ability of humans to extract regularities across experiences over time, engages the medial temporal lobe in the healthy brain. This leads to the hypothesis that statistical learning may be impaired in epilepsy patients, and that this impairment could contribute to their varied memory deficits. In turn, epilepsy patients provide a platform to advance basic understanding of statistical learning by helping to evaluate the necessity of medial temporal lobe circuitry through disease and causal perturbations. We implemented behavioral testing, volumetric analysis of the medial temporal lobe substructures, and direct electrical brain stimulation to examine statistical learning across a cohort of 61 epilepsy patients and 28 healthy controls. Behavioral performance in a statistical learning task was negatively associated with seizure frequency, irrespective of where seizures originated in the brain. The volume of hippocampal subfields CA1 and CA2/3 correlated with statistical learning performance, suggesting a more specific role of the hippocampus. Indeed, transient direct electrical stimulation of the hippocampus disrupted statistical learning. Furthermore, the relationship between statistical learning and seizure frequency was selective: behavioral performance in an episodic memory task was impacted by structural lesions in the medial temporal lobe and by antiseizure medications, but not by seizure frequency. Overall, these results suggest that statistical learning may be hippocampally dependent and that this task could serve as a clinically useful behavioral assay of seizure frequency distinct from existing neuropsychological tests. Simple and short statistical learning tasks may thus provide patient-centered endpoints for evaluating the efficacy of novel treatments in epilepsy.

## Introduction

Memory loss is a comorbidity in epilepsy, especially temporal lobe epilepsy (TLE), and has devastating consequences for quality of life.^1–3^ More than 50% of TLE patients exhibit episodic memory (EM) deficits,^3–5^ which is the ability to encode or retrieve individual autobiographical events.^6^ TLE is associated with the dysfunction of the medial temporal lobe (MTL) structures such as the hippocampus,^4, 5, 7^ which is critical for EM processing.^8, 9^ Thus, EM deficits in TLE patients may be partially explained by underlying MTL dysfunction. Indeed, numerous studies have linked EM deficits to MTL pathology.^5, 7, 10^

Epilepsy patients, including TLE, often complain of substantially greater memory disturbances in their day-to-day life than what is identified during standard neuropsychological memory evaluations.^11, 12^ Although EM deficits play a role in such complaints, anti-seizure medications (ASMs) also negatively impact memory,^13^ complicating the attribution of these deficits to epilepsy. Moreover, other memory-related functions such as spatial navigation and semantic cognition can be impaired in epilepsy.^14, 15^

Here we explore another cognitive ability, statistical learning (SL), in epilepsy patients. SL refers to the ubiquitous human ability to extract repeating patterns (or regularities) across space and time.^16, 17^ It occurs automatically, allowing for predictions of future events and adaptive behavior in new situations based on patterns learned from past experiences. SL is thought to be fundamental to the development and healthy functioning of the mind and is crucial for general human cognition, including language acquisition, object perception, spatial navigation, and conceptual knowledge.^16^

Imaging studies conducted during laboratory tasks have posited a role of MTL and hippocampus in SL.^17–20^ This is also supported by behavioral studies in patients with MTL lesions who showed SL impairment.^21, 22^ However, these were case studies with one or a small number of patients with varying etiologies other than epilepsy and damage often beyond MTL. Nevertheless, these findings suggest the novel hypothesis that SL may be impaired by epilepsy, given that it is associated with deficits in other forms of memory supported by the MTL. Here, we implemented novel methodologies to examine SL behavior in epilepsy patients. We combined these methodologies with volumetric quantification of manually segmented MTL substructures. The surgical treatment of intractable epilepsy further allowed for direct electrical brain stimulation (DES), the first targeted and reversible causal test of the necessity of the MTL for SL behavior.

Unlike in a control EM task, we found that behavioral performance in the SL task reliably predicted whether a patient’s epilepsy was under control, operationalized by their seizure frequency, regardless of where seizures originated. The volume of hippocampal subfields CA1 and CA2/3 correlated with SL performance. DES of the MTL reduced SL performance to chance. In summary, our study discovered SL deficits across epilepsy patients and showed that the MTL is critical for SL behavior. Importantly, our novel SL task provides an easily administered clinical tool with high diagnostic power to classify patients into poorly vs. well controlled epilepsy.

## Materials and methods

### Participants

A total of 89 participants were recruited to complete the SL task, including an initial 41 patients with epilepsy, 28 age-/sex-/education-matched healthy controls, and 20 additional epilepsy patients as a predictive cohort. All patients were recruited from the Yale Comprehensive Epilepsy Center. Patients without a definitive epilepsy diagnosis or with severe cognitive comorbidities were excluded from the analysis (N = 3). The initial epilepsy patient cohort (EP; N = 38) was classified based on seizure onset zone (SOZ) into TLE patients (N = 22) and extra temporal lobe epilepsy patients (ETLE; N = 16) by two clinical epileptologists based on EEG findings. As a positive control, several TLE (N = 15) and ETLE (N = 16) patients completed an additional EM task. Healthy controls (HC) completed both the SL and EM tasks. The predictive cohort (N = 20) was collected to validate how well SL could predict seizure control in new, previously unanalyzed epilepsy patients. The DES cohort (N = 5) was recruited after undergoing intracranial electroencephalography (iEEG) monitoring for seizure localization. Informed consent was given by all participants. The study was approved by the Institutional Review Board at Yale University.

### Behavioral Tasks

Two computer-based behavioral tasks were designed to evaluate SL and EM independently. The tasks were presented on a Windows laptop running a MATLAB script (R2019a; Mathworks Inc., Natick, MA) with Psychtoolbox 3.0.16.^23, 24^ SL task stimuli consisted of glyphs belonging to Sabaean and Ndyuka alphabets.^17^ EM task stimuli consisted of faces obtained from the Chicago Face Database.^25^ Participants completed the SL task, followed by the EM task.

### SL task design

The SL task was divided into two phases, an exposure phase followed by a test phase (Fig. 1a). During the exposure phase, 12 glyphs were randomly assigned into six pairs, and each pair was presented 10 times (one glyph at a time) in a continuous sequence for a total of 120 trials. The order of pairs in the sequence was random and there were no breaks or other cues between pairs. As a result, participants could only rely on the transition probabilities between glyphs to learn the pairs. Namely, the transition probability between the first and second glyph in a pair was 1.0 (i.e., the first glyph was always followed by the second glyph) whereas the transition probability between the second glyph in a pair and the first glyph of another pair was 0.2 (i.e., the second glyph in a pair was equally likely to be followed by the first glyph of the other five pairs). Each glyph was presented for 0.4 seconds followed by inter-stimulus interval of 0.8 seconds, with the full sequence lasting a total of 144 seconds.

**Figure 1.**
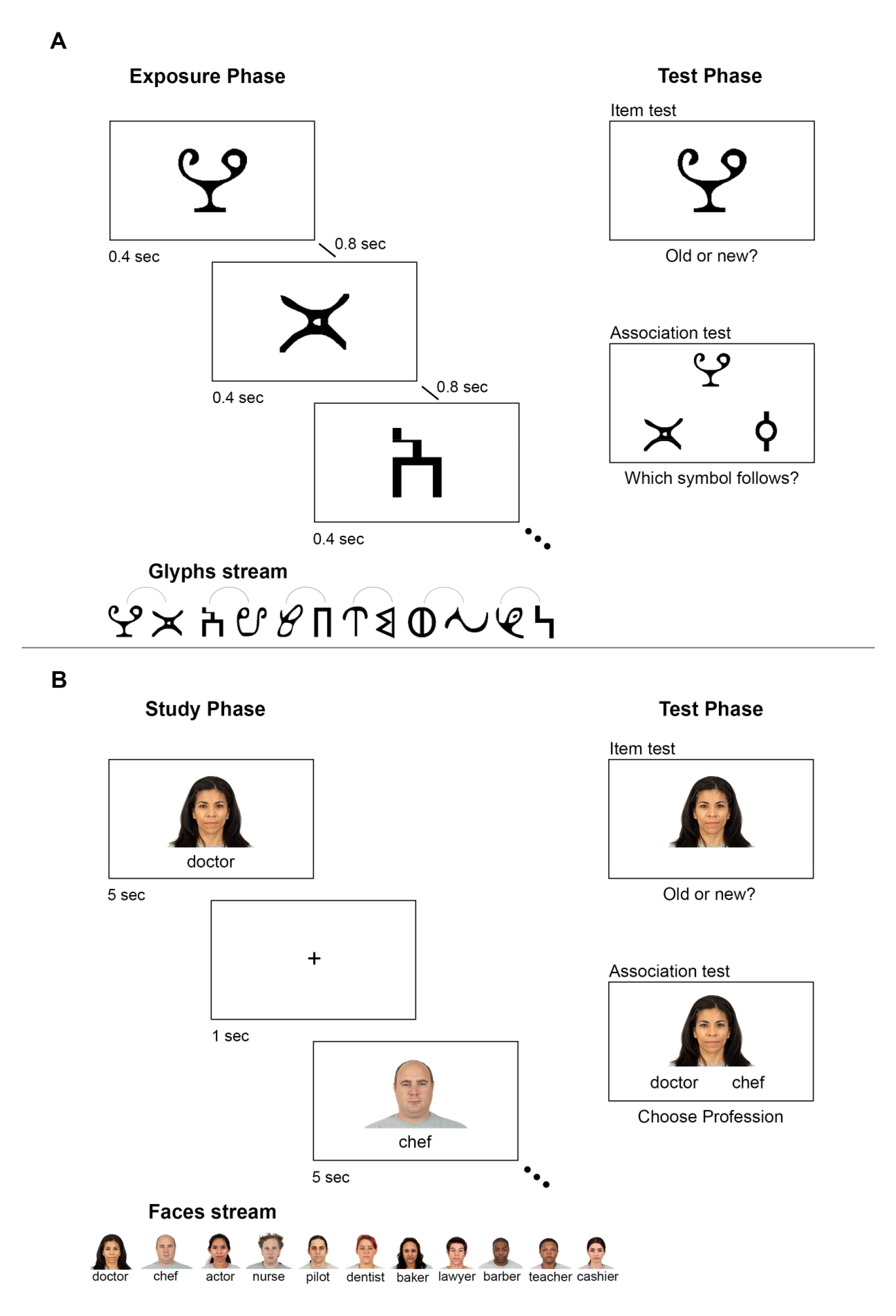
Behavioral task designs for statistical learning and episodic memory. (A) Statistical learning task. In the exposure phase, 12 glyphs were randomly assigned to pairs, presented 10 times (one glyph at a time) in a random order where the transition probability within each pair was 1.0 and between pairs was 0.2. The test phase had two parts: an item test and an association test. The association test presented the first glyph of each pair on the top of the screen and participants chose which of two glyphs was associated. (B) Episodic memory task. In study phase, face-occupation pairs were presented one pair at a time and participants were instructed to memorize each pair. In the test phase, the item test (each face was old or new) was followed by an association test (which occupation each face was associated with).

The test phase contained two parts. The first part was an item test, during which the 12 old glyphs from the exposure phase and 12 new glyphs were presented one at a time in an intermixed order for a total of 24 trials. Participants were instructed to respond whether each glyph was old (i.e., seen in the exposure phase) or new (i.e., not seen in the exposure phase). The item test assessed baseline recognition memory for the individual glyphs, which did not require statistical learning. The second part was an association test. On each trial, the first glyph of a pair was presented, and participants were asked to choose with which of two glyphs it had been paired in the exposure phase. Because both test alternatives were equally familiar items, participants needed to rely on learned transition probabilities to respond correctly. Each pair was tested twice in the association test for a total of 12 trials. The association test assessed statistical learning of the glyph pairs.

### EM task design

The EM task was divided into study and test phases, separated by a distractor task (Fig. 1b). During the study phase, 11 novel faces and 11 common occupations were grouped into pairs. Each face-occupation pair was presented once for 5 seconds followed by a 1 second inter-stimulus interval. Participants were instructed to memorize each face-occupation pair. This was followed by a distractor task during which participants solved 10 simple arithmetic problems, a standard approach in EM tasks to ensure that test performance relied on long-term memory rather than working memory (e.g., to avoid recency effects).^26^

Designed to mirror the SL task, the test phase for the EM task consisted of an item test and an association test. In the item test, participants made an old/new recognition memory judgment for 11 old faces intermixed with 11 new faces. In the association test, each studied face was presented with two studied occupations and participants needed to choose the matching profession based on the study phase. This differs from the SL task in that EM performance depended on one-shot learning of pairs presented discretely, as opposed to gradual learning of pairs embedded in a continuous sequence and extracted across multiple exposures.^27^

### MRI acquisition and MTL segmentation

For a subset of participants with epilepsy (N = 27), we manually segmented the subfields of the hippocampus and the subregions of the MTL cortex using structural MRI scans. Patients with lesions that impacted MTL segmentation were excluded from further analyses (N = 2). MRI scans were collected on a 3-T Siemens Prisma scanner with a 64-channel volume head coil. Coronal slices of turbo spin echo T2-weighted images acquired in an oblique orientation, perpendicular to the sylvian fissure (resolution = 0.52 x 0.52 mm, slice thickness = 2 mm, echo time = 90 ms, and flip angle = 150°) were used for MTL manual segmentation. In addition, magnetization prepared rapid gradient-echo (MPRAGE) T1-weighted images were used to calculate total intracranial volume (ICV). ITK-SNAP version 3.8.0^28^ was used for segmentation of the MTL from the T2 scans. We segmented hippocampal subfields subiculum, CA1, CA2/3, and dentate gyrus, and MTL cortical subregions perirhinal cortex, entorhinal cortex, and parahippocampal cortex. Segmentation was performed by a single individual, following structural landmarks of the MTL to define the borders of the regions of interest (ROIs) and cross checked by a content expert.^29–32^ For ICV computation, Freesurfer version 7.2.0 was used.^33^ Because variability in ROI volumes could be attributed to the ICV, each ROI volume was adjusted for ICV using an ANCOVA-based approach^34, 35^:

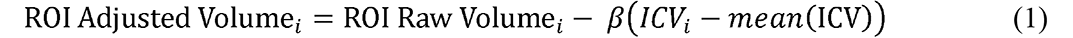

*β* is the slope of Raw Volume*_i_* (each ROI volume) regression on *ICV_i_* and *i* is the individual.

### Direct electrical brain stimulation

Five patients undergoing iEEG were recruited for DES. These patients were implanted for the clinical purpose of seizure onset localization and were selected for our study based on the location of electrodes. Electrodes were either circular 4 mm diameter subdural contacts spaced every 10 mm or 1.1 mm diameter depth electrodes with contacts spaced every 5 mm (Adtech, W.I., U.S.A.). A Nicolet stimulator (Natus, Rhode Island, U.S.A.) generated constant current, 1 Hz, biphasic square wave pulses of 300 μs per phase, at 5 mA during the exposure phase of the SL task. An epileptologist with expertise in stimulation brain mapping was present at bedside throughout the experiment to deliver the stimulation, monitor for after-discharges, and treat seizures. No stimulation was delivered during the test phase.

We first performed the SL task while stimulating the frontal pole^17, 36, 37^ to determine whether the patients showed positive behavioral evidence of SL at baseline. This was not a given because of the severity of their epilepsy, which may have impaired MTL function even before stimulation. In other words, without performing above chance prior to MTL stimulation, the hypothesized chance performance after MTL stimulation would not be interpretable. Given this lack of interpretability, and the clinical risk of seizures from DES in the MTL,^38^ we excluded two of the patients who did not show behavioral evidence of SL during frontal pole stimulation. The remaining three patients who showed SL during frontal pole stimulation received DES in the hippocampus (N = 3). A different set of glyphs was used for each region stimulated to avoid interference between task repetitions.

### Statistical analyses

We analyzed the behavioral performance of participants on the SL and EM tasks using one-sample t-tests against chance within the healthy control and epilepsy groups. Association test scores were computed as the proportion of correct responses (chance = 0.5) and item test scores were computed using d-prime, i.e., the difference between the z-transform of the hit rate and false alarm rate (chance = 0).^39^ We then compared performance across groups, split by SOZ (TLE vs. ETLE) for patients with epilepsy, and compared with HC, using analysis of variance (ANOVA) tests, followed by *post hoc* tests using Tukey’s multiple comparisons correction. The fit of the data to a normal distribution was assessed using Shapiro-Wilk test. For non-normally distributed data, a Kruskal-Wallis test was used instead of ANOVA to contrast groups, followed by *post hoc* test using Dunn’s multiple comparisons. These tests were performed using GraphPad Prism version 9 (GraphPad Software, Inc.). Significance level was set at *p* ≤ 0.05.

To test whether variables other than SOZ influenced performance, multivariate linear models (ordinary least-squares linear regressions) were used to separately predict SL and EM association test scores. Two models were run, one in which SL association test performance was the dependent variable and another in which EM association test performance was the dependent variable. The independent variables included in both models were SOZ (1 = TLE), age, sex (1 = male), ASMs known to affect memory, MTL lesions, seizure control, and item test scores from behavioral tasks. For ASMs, a binary variable was used (1 = taking ASMs that potentially affect memory). The classification of ASMs into memory-affecting and memory-sparing is presented in Supplementary Table 1. MTL lesion was based on radiological diagnosis by a clinical neuroradiologist or neurosurgeon, and a binary categorization was used to classify patients based on MTL lesion (1 = present). The degree of seizure control was defined by the seizure frequency as determined by the clinical team and confirmed by an epileptologist or neurosurgeon. A binary categorization, based on the International League Against Epilepsy (ILAE) surgical outcome classification, was employed (1 = controlled).^40^ Seizure-free patients at the time of the study or having less than four seizure days per year were classified as controlled epilepsy patients (CE; ILAE class 1-3). Patients with four or more seizure days per year were classified as uncontrolled epilepsy patients (UE; ILAE class 4-6).

A comparative analysis was performed to assess the ability of SL and EM association test scores to sort patients based on epilepsy control status. First, we ran a receiver operator characteristic (ROC) curve analysis and compared the area under the curve (AUC) for SL and EM.^41^ The parameters from these tests were then used to calculate the sample size for a new validation epilepsy cohort (power analysis with 80% confidence).^42^ This new cohort of 20 epilepsy patients performed the same tasks, and ROC and AUC computations were repeated.

To evaluate relationships between behavioral performance and MTL ROI volumes, multivariate linear regression models were employed to predict behavioral performance in SL and EM association tests. For each dependent variable, three models were run to examine different levels of granularity between MTL volumes and performance. Model A included the total bilateral hippocampal volume and total MTL cortical volume. Model B considered the anterior and posterior MTL cortical volumes, and anterior and posterior hippocampal volumes. Model C included all manually segmented hippocampal subfields and MTL cortical ROIs. All models included age and sex as control variables and volumes were adjusted for ICV. To address outliers, robust regression was used instead of least squares regression.^43^ Robust regression utilizes the leverage and residual of each data point resulting from a least square regression and assigns less weight to datapoints of large residuals, using Huber weight and Biweight, and influential points are dropped from the model.^43, 44^ For the SL association test models (where 48% of scores were ≤ 0.5), scores < 0.5 were set to chance (= 0.5); values below chance can only be interpreted as chance, and thus such variance should not meaningfully correlate with volume measurements.^45^ Both raw scores (without re-coding below chance scores as chance) and the re-coded scores were tested in all models. All regression models were computed using STATA 17 software (Stata Corporation, College Station, TX). For DES, given the small sample size of the cohort, descriptive statistics were used to report the behavioral data and the effect of stimulation.

### Data availability

All data and code used in this study are available upon reasonable request from the corresponding author.

## Results

### Participant demographics and clinical characteristics

Table 1 summarizes the demographics of the initial cohorts and the clinical profiles for all epilepsy patients. There were no significant differences in age (EP *mean* = 40.92; HC *mean* = 35.93; *U* = 451.5, *p* = 0.299; Mann-Whitney test), sex (χ^2^(1,66) = 0.330*, p* = 0.566), or education level (χ^2^(1, 61) = 0.863, *p* = 0.353; chi-square test) between healthy controls and epilepsy patients.

**Table 1.**
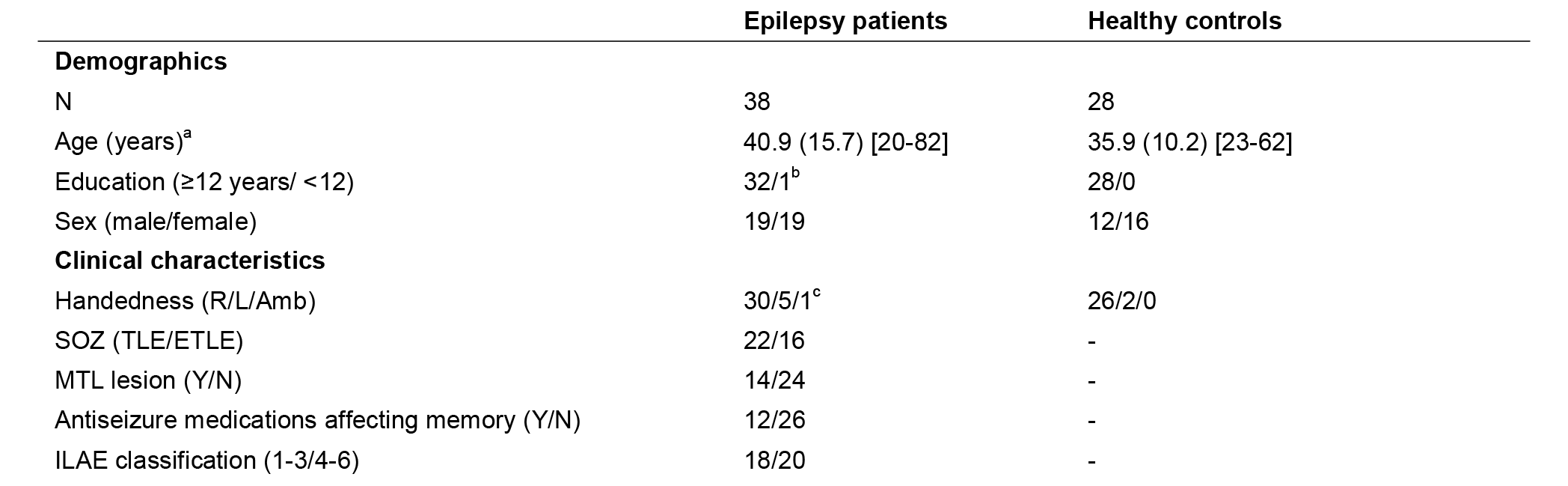
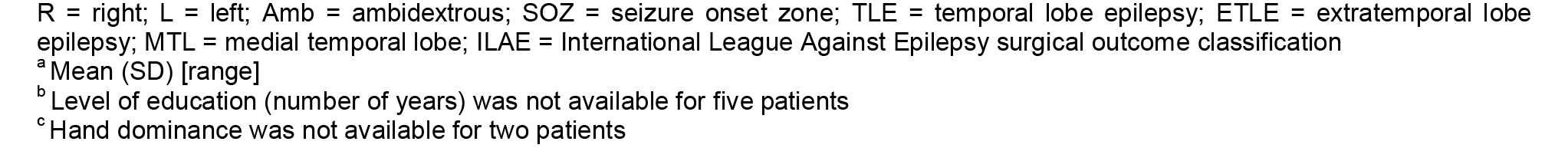
Demographics and clinical profiles

|  | Epilepsy patients | Healthy controls |
| --- | --- | --- |
| **Demographics** |  |  |
| N | 38 | 28 |
| Age (years)<sup>a</sup> | 40.9 (15.7) [20-82] | 35.9 (10.2) [23-62] |
| Education ($\geq 12$ years/ $< 12$) | 32/1<sup>b</sup> | 28/0 |
| Sex (male/female) | 19/19 | 12/16 |
| **Clinical characteristics** |  |  |
| Handedness (R/L/Amb) | 30/5/1<sup>c</sup> | 26/2/0 |
| SOZ (TLE/ETLE) | 22/16 | - |
| MTL lesion (Y/N) | 14/24 | - |
| Antiseizure medications affecting memory (Y/N) | 12/26 | - |
| ILAE classification (1-3/4-6) | 18/20 | - |

R = right; L = left; Amb = ambidextrous; SOZ = seizure onset zone; TLE = temporal lobe epilepsy; ETLE = extratemporal lobe epilepsy; MTL = medial temporal lobe; ILAE = International League Against Epilepsy surgical outcome classification
<sup>a</sup> Mean (SD) [range]
<sup>b</sup> Level of education (number of years) was not available for five patients
<sup>c</sup> Hand dominance was not available for two patients

### Behavioral results

#### Statistical learning performance

We first evaluated behavioral performance in the SL task (Fig. 2a). There was reliable SL above chance in both HC (*N* = 28, mean = 0.64, *SEM* = 0.03; t(27) = 4.816*, p* < 0.0001) and EP (*N* = 38, *mean* = 0.57, *SEM* = 0.03; *t*(37) = 2.20*, p* = 0.034) groups, which marginally differed from each other (*t*(64) = 1.77, *p* = 0.082).

**Figure 2.**
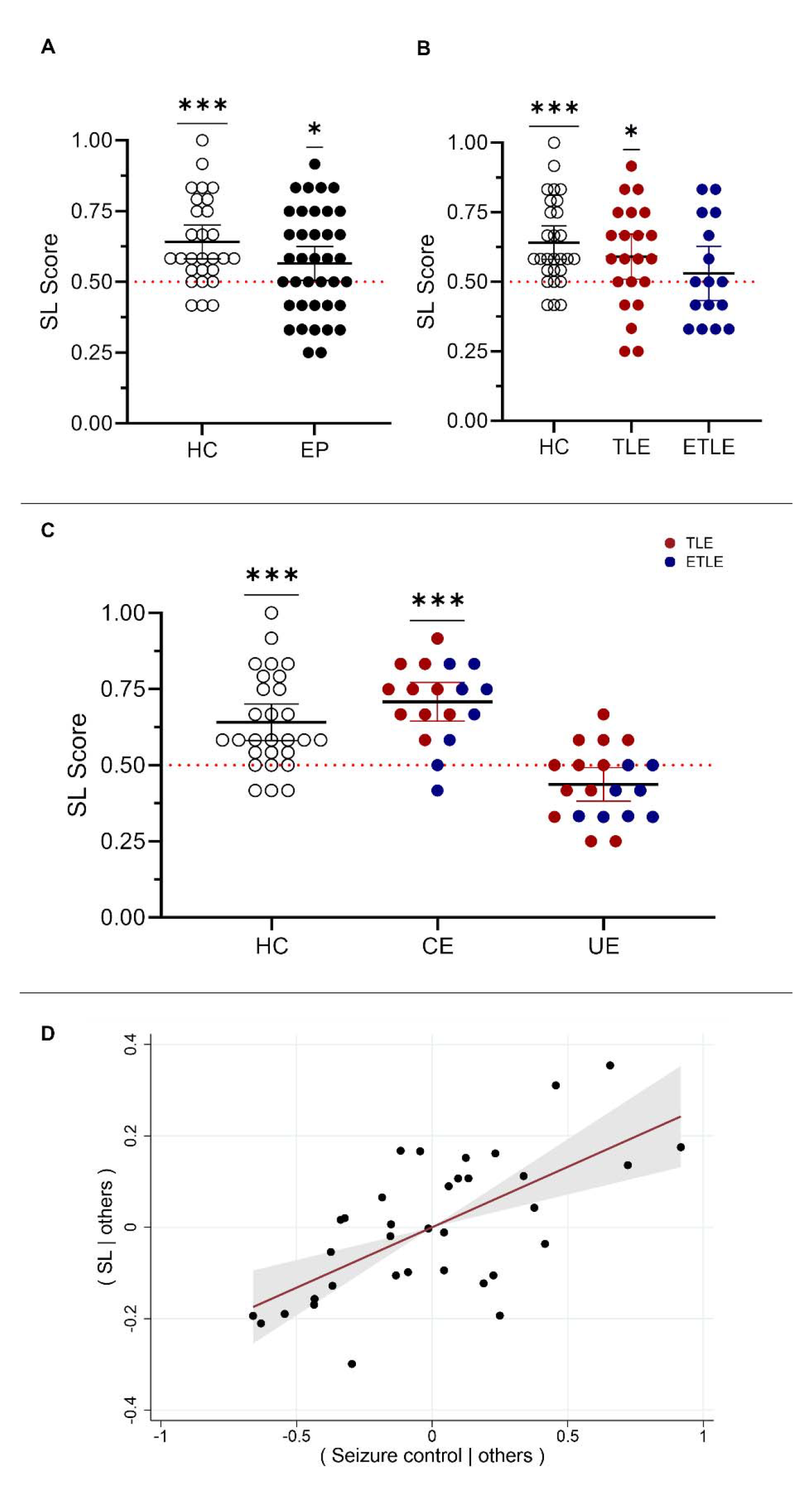
Seizure frequency predicts statistical learning performance. (A) Behavioral performance on SL task in healthy control (HC) and epilepsy patients (EP). Each dot represents an individual’s average association test score relative to chance (red dotted line, 0.5). The black lines reflects the *mean* and the error bars reflect the 95% confidence interval. (B) Dividing EP participants by seizure onset zone (TLE vs. ETLE). (C) Dividing EP participants by seizure control (controlled vs. uncontrolled epilepsy). (D) Partial relationship between seizure control and SL performance. Shading around line of best fit reflects 95% confidence interval. *** *p* < 0.001, * *p* < 0.05.

To better understand the variance in the EP group, we first divided patients based on seizure onset zone (SOZ) into TLE and ETLE subgroups and compared each subgroup to chance (Fig. 2b). The TLE subgroup performed above chance (*N* = 22, *mean* = 0.59, *SEM* = 0.04; *t*(21)= 2.323, *p* = 0.030), whereas the ETLE subgroup did not (*N* = 16, *mean* = 0.53, *SEM* = 0.05; *t*(15)= 0.665, *p* = 0.516). These differences did not manifest as a significant effect of group when comparing across HC, TLE, and ETLE (*H*(2) = 3.418, *p* = 0.181).

We then divided the EP group based on seizure control into controlled epilepsy (CE) and uncontrolled epilepsy (UE) subgroups (Fig. 2c).^40^ CE patients performed reliably above chance (CE; *N* = 18, *mean* = 0.71, *SEM* = 0.03; *t*(17) = 6.876, *p* < 0.0001), whereas UE patients did not and in fact performed slightly below chance on average (UE; *N* = 20, *mean* = 0.44, *SEM* = 0.03). Furthermore, there was a significant effect of HC, CE, and UE group on SL performance (*F*(2, 63) = 20.81*, p* < 0.0001). The UE subgroup performed worse than the CE subgroup (*p* < 0.0001; Fig. 2c) and the HC group (*p* < 0.0001).

We then built a multivariate regression model to investigate how these and other variables jointly predict SL performance in the EP group. Specifically, we evaluated SL performance relative to age, sex, ASM, SOZ, MTL lesion, item memory, and seizure control. Seizure control was the only reliable predictor of SL performance (*p* = 0.0001, partial regression coefficient = 0.264; Fig. 2d), with SOZ not reaching significance (*p* = 0.188, partial regression coefficient = 0.075). Overall, these data show that SL is impaired in patients with poorly controlled epilepsy, irrespective of SOZ.

#### Comparison to episodic memory

We next sought to replicate findings that SOZ may predict EM performance.^3–5^ Similar to SL, HC (*N* = 15, *mean* = 0.97, *SEM* = 0.02; *t*(14) = 32.39*, p* < 0.0001) and EP (*N* = 31, *mean* = 0.84, *SEM* = 0.03; *t*(30) = 11.82*, p* < 0.0001) groups performed above chance in the association test of the EM task (Fig. 3a). The EP group performed worse than the HC group (*U* = 115*, p* = 0.0032).

**Figure 3.**
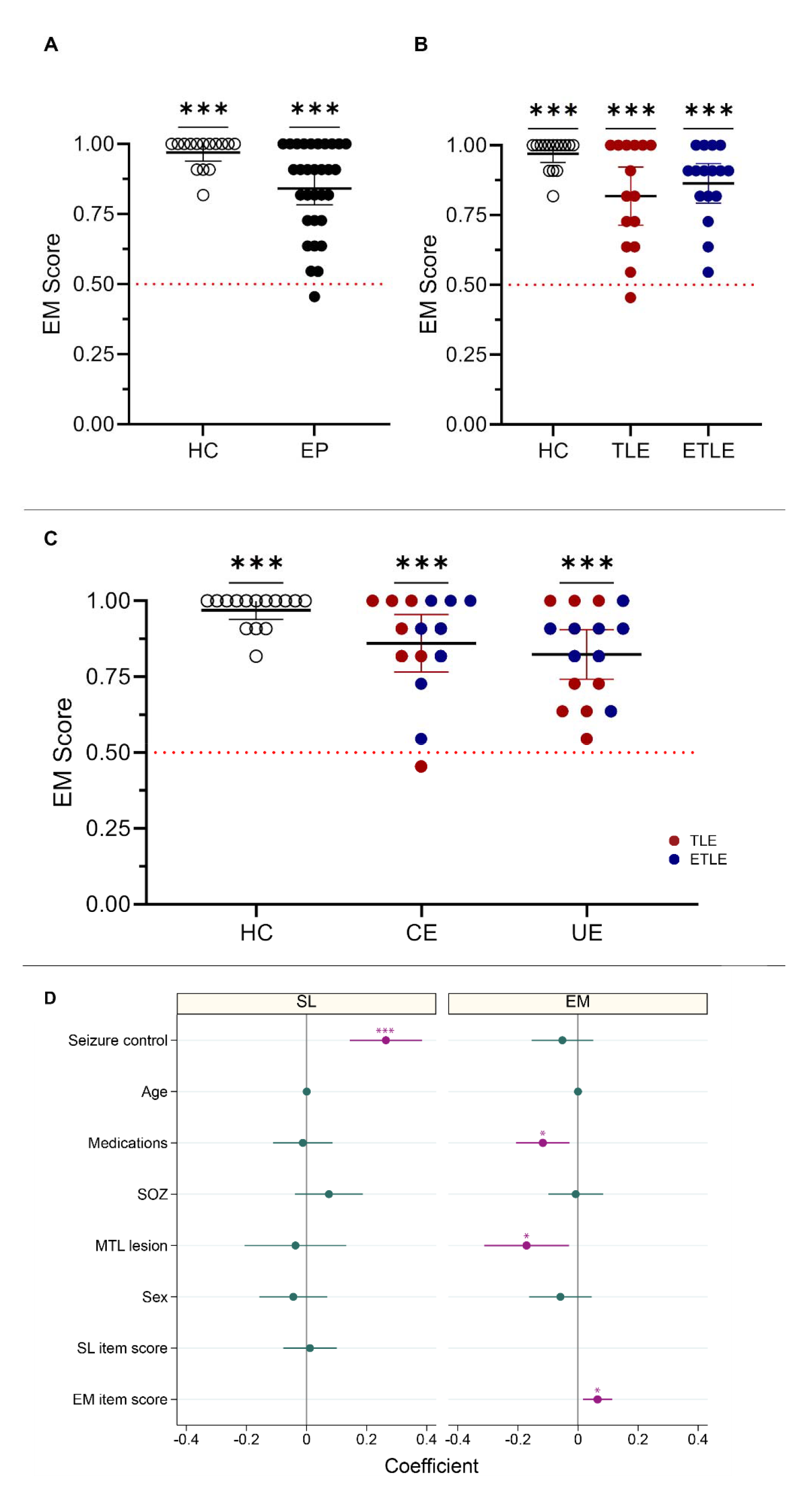
Medial temporal lobe lesion and antiseizure medications predict episodic memory performance. (A) Behavioral performance on EM task in HC and EP groups. Each dot represents an individual’s average association test score relative to chance (red dotted line, 0.5). The black lines reflect the mean and the error bars reflect 95% confidence interval. (B) Dividing EP participants by seizure onset zone (TLE vs. ETLE). (C) Dividing EP participants by seizure control (controlled vs. uncontrolled epilepsy). (D) Summary of predictors of SL and EM behavioral performance. The dot for each predictor and task reflects partial regression coefficient and the surrounding band reflects 95% confidence interval. *** *p* < 0.001. * *p* < 0.05.

The EP group was first divided based on SOZ into TLE (*N* = 15, *mean* = 0.82, *SEM* = 0.05; *t*(14) = 6.543 *p* < 0.0001) and ETLE (*N* = 16, *mean* = 0.53, *SEM* = 0.05;, *t*(15)= 10.95, *p* < 0.0001) subgroups (Fig. 3b). A three-way comparison of HC, TLE, and ETLE revealed a group effect (*H*(2) = 8.515*, p* = 0.014). Post hoc tests revealed significant differences between HC and TLE *(p* = 0.023) and HC and ETLE *(p* = 0.039), but no difference between the TLE and ETLE (*p* > 0.99).

When the EP group was divided into CE (*N* = 15, *mean* = 0.86, *SEM* = 0.04; *t*(14)= 8.414, *p* < 0.0001) and UE (*N* = 16, *mean* = 0.82, *SEM* = 0.05; *t*(15)= 8.148, *p* < 0.0001) subgroups irrespective of SOZ (Fig. 3c), we observed a group effect compared with HC (*H*(2)= 9.262*, p* = 0.0097). Post hoc tests revealed significant differences between HC and UE (*p* = 0.009), but not between HC and CE (*p* = 0.1163) or between CE and UE (*p* > 0.99).

To evaluate for predictors of EM associative performance in EP, we again ran a multivariate regression model with the same variables as the previous SL model (Fig. 3d). Poor EM performance was associated with presence of an MTL lesion (*p* = 0.018; partial regression coefficient = −0.165) and with ASMs that impact memory (*p* = 0.003; partial regression coefficient = −0.138), while EM item test score was associated with better EM associative performance (*p* = 0.009; partial regression coefficient = 0.063). In summary, whereas seizure control was the only predictor of SL associative performance, MTL lesions, ASM, and item memory predicted EM associative performance. Critically, EM did not significantly vary by seizure frequency/control.

### SL as a predictor of epilepsy severity

Given that seizure control was the strongest predictor of SL performance in epilepsy patients, we next asked whether the short behavioral SL task could be used in reverse to predict seizure frequency and epilepsy control in a new cohort of epilepsy patients. We performed an area-under-the-curve (AUC) analysis of the receiver operating characteristic (ROC) curves generated from SL and EM scores in a new cohort of patients that completed both tasks (Fig. 4). The association test scores were used as independent variables to predict seizure control as an outcome variable. SL task performance was a strong predictor of seizure control in the new cohort (AUC = 0.96), and more than EM task performance (AUC = 0.70; χ^2^(1, 19) = 3.94*, p* = 0.047).

**Figure 4.**
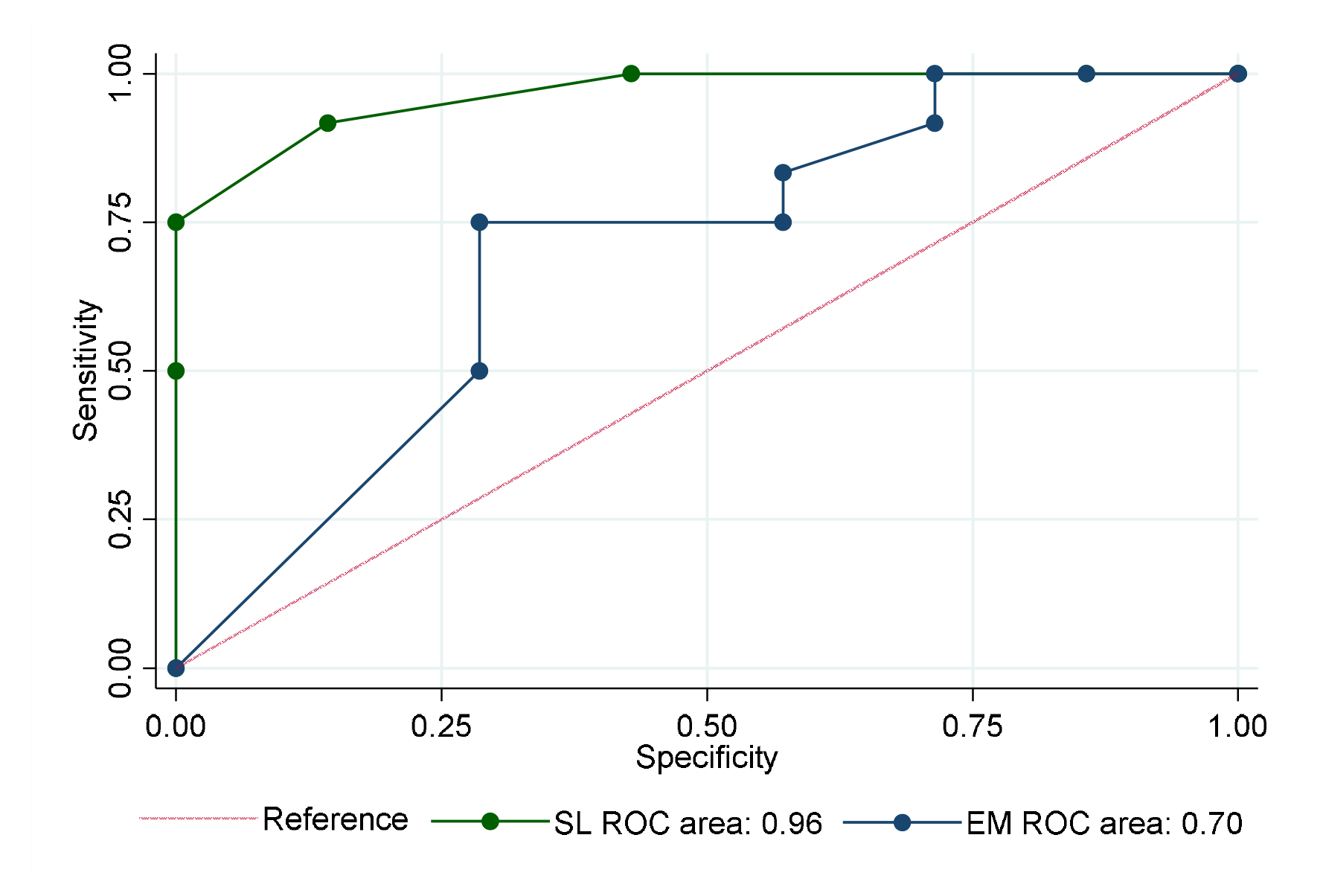
Statistical learning predicts seizure control better than episodic memory. The ability to predict seizure control differed by task. The AUCs from an ROC analysis of the SL and EM association test performance were 0.96 (95% CI: 0.90-1.00; *SEM* = 0.03) and 0.70 (95% CI:.0.43-0.96; *SEM* = 0.14), respectively.

### Relationship to MTL substructure volumes

We next built three multivariate regression models to analyze the relationship between MTL structural volumes and behavioral performance on the SL and EM association tests, examining MTL structures at different levels of granularity (each with covariates for age and sex): Model A included total hippocampal volume and total MTL cortical volume; Model B considered the anterior and posterior volumes of MTL cortex, and anterior and posterior hippocampal volumes; Model C included the volumes of hippocampal subfields subiculum, CA1, CA2/3, and dentate gyrus, and MTL subregions perirhinal cortex, entorhinal cortex, and parahippocampal gyrus (Fig. 5a). Here we report the results for the most detailed model (Model C) and the other models (Models A and B) are reported in supplementary material (Supplementary Table 2).

**Figure 5.**
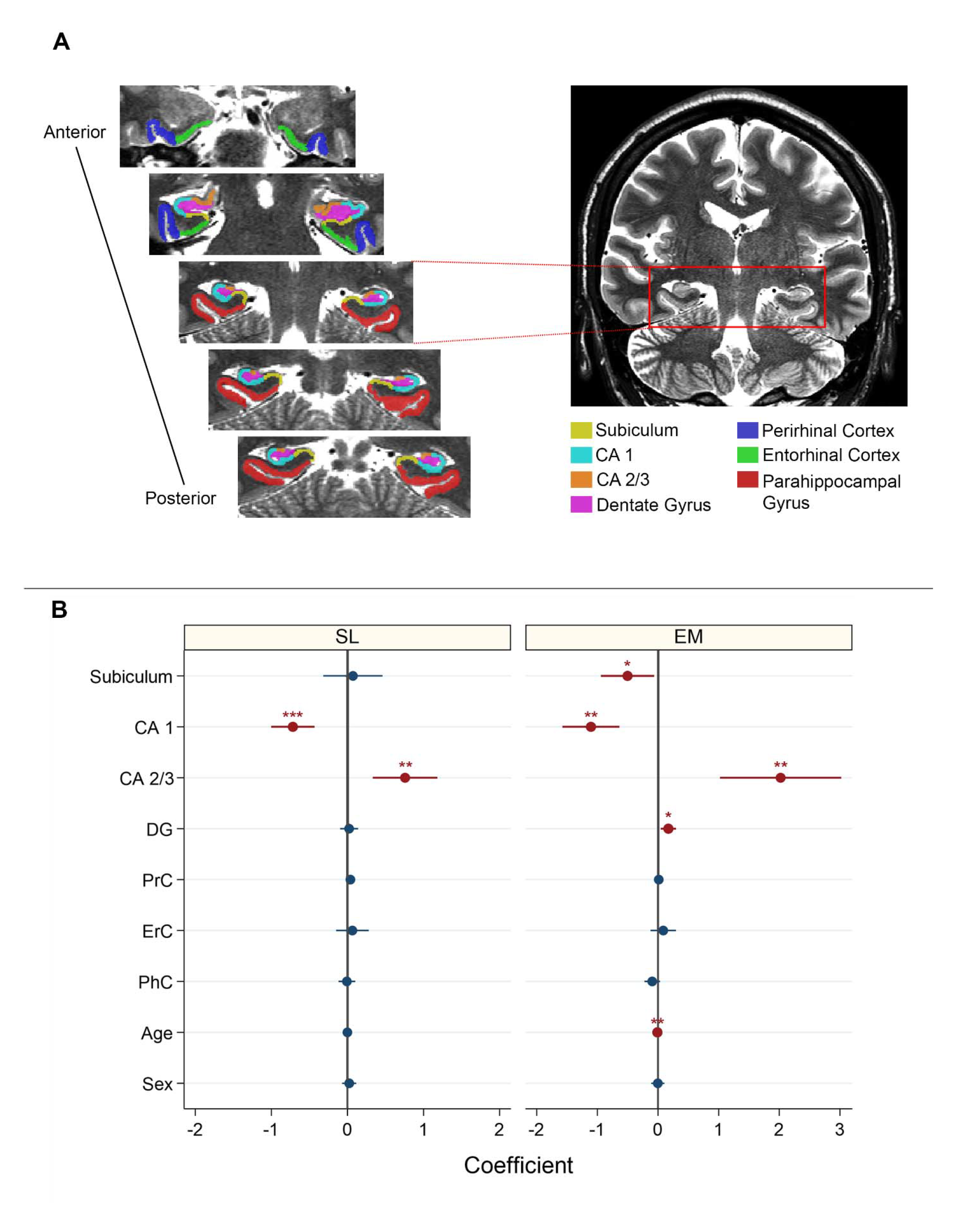
Relationship between behavioral performance and medial temporal lobe substructure volumes. (A): Manual segmentation method. Anterior slices include the perirhinal cortex (blue) and entorhinal cortex (green). Hippocampal head subfields appear along with these two cortical regions. The disappearance of the uncal apex marks the transition point between the hippocampal head and body, going posteriorly. The parahippocampal gyrus formation (red) begins with the hippocampal body. (B): Regression coefficients plots for robust multivariate regression models to predict performance in SL (left) and EM (right) association tests. *** *p* < 0.001. ** *p* < 0.01. * *p* < 0.05

The fit of Model C for SL associative performance was highly reliable overall (*N* = 25, *R*^2^ = 0.46, *F*(9, 15) = 5.34, *p* = 0.0023). The volume of hippocampal subfields CA1 (*p* = 0.0001, coefficient = −0.718) and CA2/3 (*p* = 0.002, coefficient = 0.757) reliably predicted SL performance (Fig. 5b); that is, SL performance was negatively correlated with CA1 volume and positively correlated to CA2/3 volume.

A similar pattern was observed for EM overall (*N* = 18, *R*^2^ = 0.77, *F*(9, 8) = 9.07, *p* = 0.0025) and in CA1 (*p* = 0.001, coefficient = −1.105) and CA2/3 (*p* = 0.002, coefficient = 2.022) hippocampal subfields. Additionally, subiculum volume (*p* = 0.029, coefficient = −0.502), dentate gyrus volume (*p* = 0.012, coefficient = 0.171), and patient age (*p* = 0.001, coefficient = - 0.009) were associated with EM performance (Fig. 5b).

### Stimulation experiment

Given that the volumes of MTL subregions were associated with SL performance, we next asked whether the MTL is causally necessary for SL.^21, 22^ We used transient DES in a new cohort of patients to disrupt hippocampal function during the exposure phase of the SL task (Fig. 6a). We performed MTL stimulation in the three patients who exhibited SL during baseline stimulation of the frontal pole as a positive control (scores = 0.67, 0.83, 0.67). In contrast, when we stimulated the hippocampus in these patients (Fig. 6b), they no longer showed learning (scores = 0.50, 0.59, 0.50, respectively; Fig. 6c; Supplementary Table 3).

**Figure 6.**
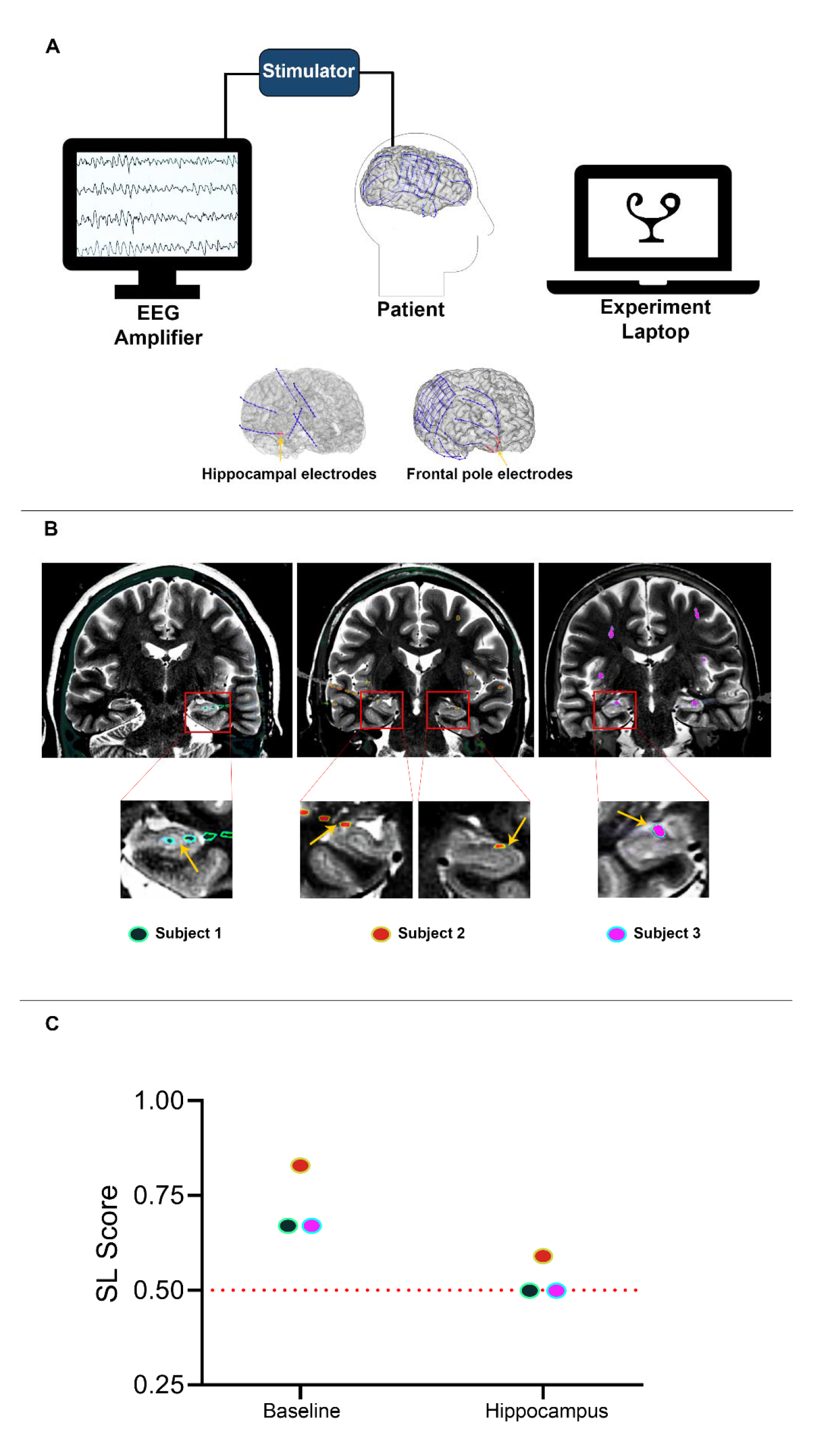
Necessity of the hippocampus for statistical learning. (A): Schematic diagram of the stimulation experiment. Three patients performed the SL task twice. DES was administered at 1 Hz during the exposure phase of the task (Fig. 1a; only change from prior experiments was that each glyph was probed once rather than twice in the association test) through a cortical stimulator connected to the patient’s leads. Example of electrode localization for the contacts representing hippocampus and frontal pole of one patient (generated using Freesurfer and iElectrodes software with pre-op MRI and post-op CT scans) shown at the bottom. (B): Electrode localization for each of the three subjects included in the stimulation experiment (electrodes shown on CT scan overlaid on T2-weighted MRI). Two patients who did not show SL during baseline frontal pole stimulation were excluded. Distinct colors are applied for each patient’s contacts. Yellow arrows indicate where the stimulation was administered (bipolar stimulation). (C): SL scores for three patients in the SL association test when the control region (frontal pole) vs. hippocampus were stimulated. The red-dotted line represents chance performance (0.5). Stimulation of the hippocampus eliminated the positive evidence of SL observed during the baseline stimulation of the control region. Colors correspond to patients as indicated in (B).

## Discussion

In the current study, we investigated how statistical learning (SL) — a fundamental form of human learning which is used to acquire the structure of our environment, make predictions, and behave efficiently — is impacted by epilepsy. We were motivated by recent demonstrations that SL depends on the medial temporal lobe (MTL) ^21, 22^ and that epilepsy can be associated with MTL dysfunction and memory deficits.^4, 5, 7^ We found that SL is impaired in patients with uncontrolled epilepsy, irrespective of where their seizures originated. Using a new cohort, we validated that the SL task can reliably predict if seizures are controlled in epilepsy patients. Furthermore, manual segmentation and quantification of MTL volumes in epilepsy patients showed that SL performance was related to the volumes of hippocampal subfields CA1 and CA2/3. Finally, in a subset of intracranial patients with normal SL during baseline frontal stimulation, DES of the hippocampus during the exposure phase of the SL task impaired associative performance. Taken together, our results suggest that poorly controlled epilepsy may impair SL via disruption of the MTL (including the hippocampus). In addition, SL task performance may be a clinically relevant tool to evaluate seizure frequency and severity in epilepsy patients. The findings in our EM task are concordant with cumulative evidence suggesting that the MTL is critical for EM.^8, 9^

Given prior findings that SL may rely on the MTL,^21^ we hypothesized that SL would be impaired in TLE. However, in our study, the location of the SOZ did not impact SL task performance. Rather, seizure control — frequency of seizure days per year — reliably predicted SL performance (and vice versa in the validation cohort) for epilepsy patients including both TLE and ETLE. One possible reason is that repetitive seizures (regardless of onset) may lead to structural damage or functional impairment in the MTL and other brain regions relevant for SL.^46–48^ Our findings may thus be consistent with a previous report of impairments in auditory SL in a cohort of patients with multi-lobar brain damage from strokes.^49^

We did not find the same relationship between seizure control and EM. Although we may have expected this relationship, there have been conflicting results on the relationship between seizure frequency and declarative memory.^50–53^ In our study, some patients who exhibited normal EM performance showed chance level performance in the SL test. This raises the question as to whether the differential effect of seizure frequency on these two behaviors depend on the networks disrupted by seizures, and whether these broad networks differentially support SL and EM.

After manually segmenting and quantifying MTL substructure volumes, we found positive relationships between hippocampal subfield CA2/3 volume and performance on both the SL and EM tasks. The volume-EM relationship observed here is in line with prior work on cell counts and histopathology analysis of resected hippocampal tissue in epilepsy patients, for example showing that neuronal loss in hippocampal subfields (excluding CA1) correlates with declarative memory performance.^54^ SL performance was also positively correlated with CA2/3 volumes, suggesting involvement of the hippocampus in SL. However, the positive correlation of CA2/3 volume with SL performance observed here is inconsistent with a previous study that found a negative correlation between CA2/3 volume (in the hippocampal head) and SL.^35^ A potential explanation for this difference could be the difference in age between the two samples (our sample was older) and, more importantly, the pathology (we only considered epilepsy patients for the volumetric analysis, whereas they considered healthy individuals).^35^ Indeed, epilepsy (especially TLE) is often associated with structural disease.^55^ The negative correlation we observed between CA1 volume and performance in SL and EM is in line with histological studies in hippocampal sclerosis (HS) ILAE type 2 patients, i.e., patients with predominantly CA1 neuronal cell loss, who do not exhibit declarative memory impairment.^54, 56, 57^ Furthermore, a negative relationship between EM performance and CA1 volume (compared to the other subfields, measured by cell loss) has previously been reported in epilepsy.^58^

Finally, we performed transient direct electrical stimulation to test for functional role of the MTL in SL. We found that hippocampal stimulation during the exposure phase of the SL task disrupted performance relative to baseline frontal stimulation. This finding suggests that the MTL may be causally involved in SL. Our findings support previous fMRI and lesion-based studies that implicated the MTL in SL.^17–22^ However, given our limited sample size, we suggest caution in the interpretation of this generalization of our results. Notably, while our DES protocol allows us to test whether the MTL is necessary for SL, it does not evaluate whether frontal pole is necessary (we excluded patients who did not learn during frontal pole stimulation, because this was our non-MTL positive control). We also note a limitation of our stimulation procedure: we always stimulated the hippocampus last because of its highly epileptogenic nature.^59^ This is particularly relevant because SL can be affected by order: exposure to one set of regularities may block or interfere with the subsequent learning of a second set.^60–62^ We accepted these compromises in experimental design given the rare opportunity to perform reversible causal perturbation of the human brain, and we acknowledge the resulting limitations on what can be concluded from the findings. Although future work is needed to verify and expand upon these findings, we believe that they provide supportive evidence for a causal link between the MTL, specifically the hippocampus, and SL.

Beyond providing a novel assessment of cognition in epilepsy patients, a potentially impactful conclusion of our study is that the SL task can be used to sort epilepsy patients based on seizure frequency (i.e., controlled vs. uncontrolled epilepsy). Notably, the EM task did not have this predictive power. This is clinically relevant as most neuropsychological testing batteries are based on EM tasks for the memory assessment section and thus may not capture the full range of memory decline or MTL dysfunction in epilepsy.^3, 63^ We believe that a short but powerful task such as the SL task may provide added value in classifying seizure frequency, as it might surpass the current methodology of seizure diaries, which are unreliable in documenting seizure frequency.^64^ Thus, the SL task has potential for both seizure severity assessment and comprehensive cognitive assessment. The finding that patients can perform well in EM and poorly at SL is notable as it may explain some of the subjective memory-related complaints of patients with epilepsy that are not reflected on the current objective clinical testing measures.^11, 14, 15^ Therefore, adding the SL task to the current neuropsychological assessment protocol as part of the clinical evaluation may expand patients’ cognitive assessment. We believe that our findings motivate the consideration of SL as an important aspect in the evaluation of epilepsy patients regardless of the type of epilepsy.

In conclusion, by combining behavior, neuroanatomy, and direct electrical stimulation, our study provided a comprehensive initial investigation of how SL is affected by epilepsy. In demonstrating a novel link between SL deficit and seizure frequency, we highlight the potential utility of studying SL in epilepsy. First, characterizing SL deficits in epilepsy may lead to a more comprehensive understanding of the memory problems associated with epilepsy. Second, SL tasks have potential as a clinically relevant tool for assessing epilepsy severity.

## Abbreviations

ASM: antiseizure medication
CA: cornu ammonis
CE: controlled epilepsy
DES: direct electrical stimulation
EM: episodic memory
ETLE: extratemporal lobe epilepsy
ICV: intracranial volume
MTL: medial temporal lobe
ROI: region of interest
SL: statistical learning
TLE: temporal lobe epilepsy
UE: uncontrolled epilepsy

## Acknowledgments

We thank all members of Yale Comprehensive Epilepsy Center for providing exceptional patient care and all patients that participated in the study. We also thank Joseph King and Layton Lamsam for fruitful discussions on earlier versions of this work.

## Supplementary material

Separate document

