## Supplementary for "Statistical learning in epilepsy: Behavioral, anatomical, and causal mechanisms in the human brain"

**Supplementary Table 1 Antiseizure medications taken by the cohort of epileptic patients and their impact on memory**

| | Epilepsy patients n | | | Impact on memory | Reference |
|---|---|---|---|---|---|
| **ASM type** | **All** | **Controlled** | **Uncontrolled** | | |
| Phenytoin (Dilantin)[1] | 3 | 1 | 2 | ↓ | Aldenkamp 2002 |
| Carbamazepine (Tegretol)[2] | 4 | 2 | 2 | ↓ | Witt et al. 2013 |
| Clobazam (Onfi)[2] | 9 | 3 | 6 | None | Witt et al. 2013 |
| Zonisamide (Zonegran)[3] | 3 | 2 | 1 | ↓ | Park et al. 2007 |
| Pregabalin (Lyrica)[4] | 5 | 0 | 5 | None | Ciesielski et al. 2006 |
| Topiramate (Topamax)[5] | 5 | 3 | 2 | ↓ | Ramsay et al. 2008 |
| Lamotrigine (Lamictal)[2] | 9 | 5 | 4 | None | Witt et al. 2013 |
| Oxcarbazepine (Trileptal)[2] | 6 | 2 | 4 | None | Witt et al. 2013 |
| Levetiracetam (Keppra)[6] | 18 | 10 | 8 | None | Gomer et al. 2007 |
| Valproic acid[7] | 1 | 1 | 0 | ↓ | Ortinski et al. 2004 |
| Lacosamide (Vimpat)[8] | 8 | 3 | 5 | None | Lancman et al. 2016 |
| Clonazepam | 3 | 0 | 3 | Unknown | |
| Lorazepam (Ativan)[9] | 2 | 0 | 2 | ↓ | Satzger et al. 1990 |
| Eslicarbazepine acetate (Aptiom)[10] | 2 | 0 | 2 | None | Milovan et al. 2010 |
| Cenobamate (Xcopri) | 3 | 0 | 3 | Unknown | |
| Cannabidiol (Epidiolex)[11] | 2 | 0 | 2 | None | Martin et al. 2019 |
| Sodium valproate (depakote)[7] | 2 | 0 | 2 | ↓ | Ortinski et al. 2004 |
| Perampanel (Fycompa)[12] | 1 | 0 | 1 | None | Meador et al. 2016 |
| Brivaracetam (Briviact)[13] | 2 | 0 | 2 | None | Biton et al. 2013 |
| Citalopram | 1 | 0 | 1 | Unknown | |
| Midazolam[14] | 1 | 0 | 1 | ↓ | Riss et al. 2008 |
| Methylphenidate (Concerta)[15] | 1 | 0 | 1 | Unknown | Adams et al. 2017 |

ASM = antiseizure medication; ↓ = potential negative impact on memory; None = no deficits

**Supplementary Table 2 Volume-behavior multivariate regression models**

| | SL associative | | EM associative | | SL item | | EM item | |
|---|---|---|---|---|---|---|---|---|
| | **P-value** | **Coefficient** | **P-value** | **Coefficient** | **P-value** | **Coefficient** | **P-value** | **Coefficient** |
| **Model A**[1] | | | | | | | | |
| Hippocampus | 0.651 | -0.024 | 0.254 | -0.075 | 0.825 | 0.042 | 0.119 | -0.522 |
| MTL cortex | 0.458 | 0.017 | 0.689 | 0.010 | 0.366 | 0.073 | 0.975 | 0.004 |
| Age | 0.689 | -0.001 | **0.033** | **-0.008** | 0.178 | -0.014 | 0.365 | -0.015 |
| Sex | 0.731 | -0.025 | 0.228 | -0.105 | 0.741 | -0.088 | 0.260 | -0.487 |
| **Model B**[2] | | | | | | | | |
| Anterior MTL cortex | 0.136 | 0.048 | 0.595 | 0.034 | 0.956 | -0.006 | **0.003** | **0.479** |
| Posterior MTL cortex | 0.310 | -0.084 | 0.679 | -0.066 | 0.393 | 0.249 | **0.004** | **-1.141** |
| Anterior hippocampus | 0.957 | -0.004 | 0.835 | 0.022 | 0.133 | -0.447 | **0.041** | **-0.494** |
| Posterior hippocampus | 0.632 | 0.062 | 0.664 | -0.133 | 0.263 | 0.613 | **0.026** | **1.553** |
| Age | 0.520 | -0.002 | 0.053 | -0.009 | 0.951 | -0.001 | 0.496 | -0.006 |
| Sex | 0.707 | -0.026 | 0.153 | -0.133 | 0.620 | -0.110 | 0.074 | -0.350 |
| **Model C**[3] | | | | | | | | |
| Entorhinal cortex | 0.528 | 0.064 | 0.368 | 0.087 | **< 0.0001** | **0.938** | **0.0003** | **-1.555** |
| Perirhinal cortex | 0.124 | 0.038 | 0.607 | 0.013 | 0.973 | 0.001 | **0.001** | **0.374** |
| Parahippocampal cortex | 0.879 | 0.008 | 0.125 | -0.065 | 0.196 | -0.084 | 0.060 | 0.286 |
| CA1 | **0.0001** | **-0.718** | **0.001** | **-1.105** | 0.920 | -0.017 | **< 0.0001** | **-7.131** |
| CA2/3 | **0.002** | **0.757** | **0.002** | **2.022** | **0.0003** | **-1.181** | **< 0.0001** | **12.62** |
| Subiculum | 0.699 | 0.071 | **0.029** | **-0.502** | **0.002** | **1.006** | 0.163 | 0.821 |
| Dentate Gyrus | 0.703 | 0.022 | **0.012** | **0.171** | 0.068 | 0.155 | 0.516 | 0.097 |
| Age | 0.600 | -0.001 | **0.001** | **-0.009** | **< 0.0001** | **-0.027** | **< 0.0001** | **-0.054** |
| Sex | 0.611 | 0.023 | 0.947 | -0.003 | 0.499 | -0.039 | **0.0001** | **1.155** |

MTL = medial temporal lobe; HPC = hippocampus.
[1]Model A: SL associative: N = 25; $R^2$ = 0.05, F(4, 20) = 0.26, P = 0.90; EM associative: N = 19, $R^2$ = 0.35, F(4, 14) = 1.65, P = 0.216; SL item: N = 22, $R^2$ = 0.21, F(4, 17) = 1.20, P = 0.35; EM item: N = 19, $R^2$ = 0.14, F(4, 14) = 1.02, P = 0.43
[2]Model B: SL associative: N = 25 $R^2$ = 0.14, F(6, 18) = 0.4 9 , P = 0.81; EM associative: N = 19, $R^2$ = 0.41, F(6, 12) = 1.13 , P = 0.40; SL item: N = 22, $R^2$ = 0.38, F(6, 15) = 1.74 , P = 0.18; EM item: N = 19, $R^2$ = 0.39, F(6, 12) = 4.25 , P = 0.0159
[3]Model C: SL associative: N = 25, $R^2$ = 0.46, F(9, 15) = 5.34, P = 0.0023; EM associative: N = 18, $R^2$ = 0.77, F(9,8) = 9.07, P = 0.0025; SL item: N = 22, $R^2$ = 0.60, F(9,12) = 55.47, P < 0.0001; EM item: N = 18 $R^2$ = 0.65, F(9, 8) = 21.73, P = 0.0001
Bold highlights statistically significant values.

**Supplementary Table 3 SL performance in patients receiving 1 Hz stimulation by site**

| Subject id | Baseline stimulation[a] | | Hippocampus | | No stimulation[c] | | Amperage | | Hemisphere |
|---|---|---|---|---|---|---|---|---|---|
| | Assoc | Item[b] | Assoc | item | Assoc | item | min 1 | min 2 | |
| 1 | 0.67 | 3.11 | 0.50 | 1.64 | 0.67 | 3.46 | 5 mA | 10 mA | L |
| 2 | 0.83 | 2.16 | 0.59[d] | 2.46 | 0.67 | 3.12 | 5 mA | 10 mA | L & R |
| 3 | 0.67 | 2.77 | 0.50 | 2.77 | - | - | 1 mA | 5 mA | R |

min = minute; L = left; R = right
[a]Frontal pole was the control region.
[b]item scores are measured by d'
[c]Refers to behavioral performance when patients were tested in the SL task outside of surgical stay.
[d]Average of two scores (left and right hippocampal stimulation)
